# Robust antiviral humoral immunity induced by JN.1 monovalent mRNA vaccines against a broad range of SARS-CoV-2 Omicron subvariants including JN.1, KP.3.1.1 and XEC

**DOI:** 10.1101/2024.11.20.624471

**Authors:** Keiya Uriu, Yu Kaku, Yoshifumi Uwamino, Hiroshi Fujiwara, Fumitake Saito, The Genotype to Phenotype Japan (G2P-Japan) Consortium, Kei Sato

## Abstract

As of November 2024, SARS-CoV-2 Omicron JN.1 subvariants, such as KP.2 (JN.1.11.1.2), KP.3 (JN.1.11.1.3), KP.3.1.1 (JN.1.11.1.3.1.1), and XEC — a recombinant lineage between KS.1.1 (JN.13.1.1.1) and KP.3.3 (JN.1.11.1.3.3) — have been circulating in several countries. To control the infection with SARS-CoV-2 Omicron JN.1 subvariants, JN.1 monovalent mRNA vaccines have been developed. Some previous reports showed that the JN.1 monovalent mRNA vaccine of Pfizer/BioNTech (US/Germany) increased antiviral humoral immunity against JN.1 subvariants and XEC. However, the efficacy of other available JN.1 monovalent mRNA vaccines (e.g., Daiichi-Sankyo, Japan) remains unassessed. To validate the antiviral efficacy induced by JN.1 mRNA vaccines, sera were collected from individuals vaccinated with Pfizer/BioNTech JN.1 mRNA vaccine (N=15) or Daiichi-Sankyo JN.1 mRNA vaccine (N=19) before and 3-4 weeks after vaccination. We then performed a neutralization assay using these sera and pseudoviruses. Both Pfizer/BioNTech JN.1 vaccine (2.4-to 8.0-fold, P=0.0001) and Daiichi-Sankyo JN.1 vaccine (2.3-to 13-fold, P=0.0001) boosted antiviral humoral immunity against all variants tested with statistical significance. While the Pfizer/BioNTech mRNA vaccine encodes the full-length JN.1 spike (S), the Daiichi-Sankyo mRNA vaccine encodes the receptor-binding domain of JN.1 S. Our data suggest that the receptor-binding domain of JN.1 S can effectively induce antiviral humoral immunity against JN.1 subvariants and XEC comparable to the full-length JN.1 S. However, it should be considered that the sizes of our cohorts are relatively small (<20 donors per cohort), and donor characteristics, such as age, sex, underlying disease status, and previous SARS-CoV-2 infection, may critically affect the experimental results. Future investigations with larger cohorts will address this concern. When compared to vaccination with JN.1 mRNA vaccines, our previous investigations showed that the natural infection of JN.1 and KP.3.3 elicited poorer antiviral humoral immunity against JN.1 and its subvariants. Our results suggest that the JN.1 mRNA vaccination more robustly induces antiviral humoral immunity against recent JN.1 subvariants than the natural infection of JN.1 subvariants regardless of manufacturer. Moreover, as we reported last year, the humoral immunity induced by XBB.1.5 monovalent mRNA vaccine against XBB.1.5 was weaker than that against ancestral B.1.1. However, in the case of JN.1 monovalent mRNA vaccine, here we showed that the 50% neutralization titer against XBB.1.5 is greater than that against ancestral B.1.1. These observations imply that immune imprinting has shifted from that biased toward pre-Omicron to that biased toward Omicron, depending on the time and/or number of immune stimuli (e.g., infection and/or vaccination).

## Text

As of November 2024, SARS-CoV-2 Omicron JN.1 subvariants, such as KP.2 (JN.1.11.1.2)^1^, KP.3 (JN.1.11.1.3)^2^, KP.3.1.1 (JN.1.11.1.3.1.1)^3^, and XEC^4^ — a recombinant lineage between KS.1.1 (JN.13.1.1.1) and KP.3.3 (JN.1.11.1.3.3) — have been circulating in several countries. To control the infection with SARS-CoV-2 Omicron JN.1 subvariants, JN.1 monovalent mRNA vaccines have been developed. Some previous reports showed that the JN.1 monovalent mRNA vaccine of Pfizer/BioNTech (US/Germany) increased antiviral humoral immunity against JN.1 subvariants and XEC.^5-7^ However, the efficacy of other available JN.1 monovalent mRNA vaccines (e.g., Daiichi-Sankyo, Japan) remains unassessed.

To validate the antiviral efficacy induced by JN.1 mRNA vaccines, sera were collected from individuals vaccinated with Pfizer/BioNTech JN.1 mRNA vaccine (N=15; **Figure A**) or Daiichi-Sankyo JN.1 mRNA vaccine (N=19; **Figure B**) before and 3-4 weeks after vaccination. We then performed a neutralization assay using these sera and pseudoviruses. Both Pfizer/BioNTech JN.1 vaccine (2.4-to 8.0-fold, *P*=0.0001) (**Figure A**) and Daiichi-Sankyo JN.1 vaccine (2.3-to 13-fold, *P*=0.0001) (**Figure B**) boosted antiviral humoral immunity against all variants tested with statistical significance. While the Pfizer/BioNTech mRNA vaccine encodes the full-length JN.1 spike (S), the Daiichi-Sankyo mRNA vaccine encodes the receptor-binding domain of JN.1 S. Our data suggest that the receptor-binding domain of JN.1 S can effectively induce antiviral humoral immunity against JN.1 subvariants and XEC comparable to the full-length JN.1 S. However, it should be considered that the sizes of our cohorts are relatively small (<20 donors per cohort), and donor characteristics, such as age, sex, underlying disease status, and previous SARS-CoV-2 infection, may critically affect the experimental results. Future investigations with larger cohorts will address this concern.

**Figure.**
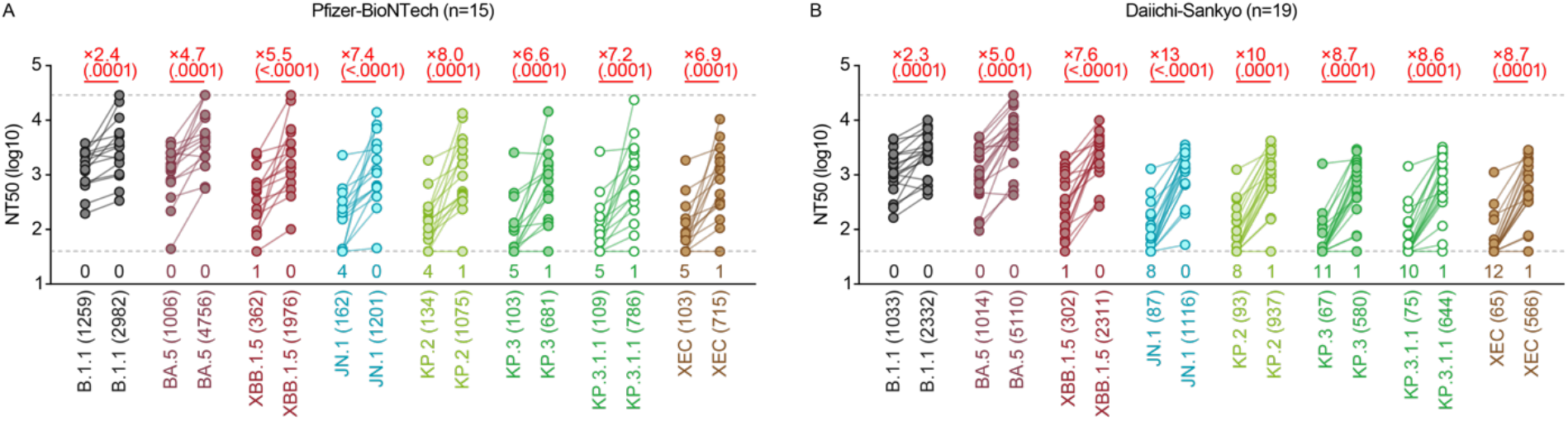
The neutralization activity induced by JN.1 monovalent mRNA vaccines. Neutralization assay. Assays were performed with pseudoviruses harboring the spike proteins of B.1.1, BA.5, XBB.1.5, JN.1, KP.2, KP.3, KP.3.1.1 and XEC. The following JN.1 monovalent mRNA vaccine sera from fully vaccinated individuals who had received Pfizer-BioNTech JN.1 vaccine (“Pfizer-BioNTech” cohort) (two 2-dose vaccinated, two 3-dose vaccinated, five 4-dose vaccinated, three 5-dose vaccinated and three 7-dose vaccinated; fifteen donors, average age: 42.1, range: 27–85, 66.7% male) (**A**) and those from fully vaccinated individuals who received Daiichi-Sankyo JN.1 vaccine (“Daiichi-Sankyo” cohort) (two 2-dose vaccinated, one 4-dose vaccinated, one 5-dose vaccinated, four 6-dose vaccinated and eleven 7-dose vaccinated; nineteen donors, average age: 57.8, range: 25–84, 26.3% male) (**B**). Assays for each serum sample were performed in quadruplicate to determine the 50% neutralization titer (NT_50_). Each dot represents one NT_50_ value of each donor, and the NT_50_ values for the same donor before and after vaccination are connected by a line. Numbers in parentheses below the graphs indicate the geometric mean of the NT_50_ value. The horizontal dashed lines indicate the detection limit of the lowest (40-fold dilution) and the highest (29,160-fold dilution), respectively. The number of sera with the NT_50_ values below the lower detection limit is shown in the figure (under the bottom horizontal dashed line). Neutralization titers below the lower and above the higher detection limit were calculated as titers of 40 and 29,160. The fold changes of NT_50_ between before and after vaccination indicated above the graph with “x” are calculated as the geometric mean of the ratio of NT_50_ obtained from each individual. The p-values in the parentheses were determined by two-sided Wilcoxon signed-rank tests.

When compared to vaccination with JN.1 mRNA vaccines (**Figure**), our previous investigations showed that the natural infection of JN.1 and KP.3.3 elicited poorer antiviral humoral immunity against JN.1 and its subvariants.^1-4^ Our results suggest that the JN.1 mRNA vaccination more robustly induces antiviral humoral immunity against recent JN.1 subvariants than the natural infection of JN.1 subvariants regardless of manufacturer.

Moreover, as we reported last year, the humoral immunity induced by XBB.1.5 monovalent mRNA vaccine against XBB.1.5 was weaker than that against ancestral B.1.1.^8^ However, in the case of JN.1 monovalent mRNA vaccine, here we showed that the 50% neutralization titer against XBB.1.5 is greater than that against ancestral B.1.1 (**Figure**). These observations imply that immune imprinting has shifted from that biased toward pre-Omicron to that biased toward Omicron, depending on the time and/or number of immune stimuli (e.g., infection and/or vaccination).

## Grants

Supported in part by AMED ASPIRE Program (JP24jf0126002, to G2P-Japan Consortium and Kei Sato); AMED SCARDA Japan Initiative for World-leading Vaccine Research and Development Centers “UTOPIA” (JP243fa627001h0003, to Kei Sato); AMED SCARDA Program on R&D of new generation vaccine including new modality application (JP243fa727002, to Kei Sato); AMED Research Program on Emerging and Re-emerging Infectious Diseases (23fk0108583, JP24fk0108690, to Kei Sato); JSPS KAKENHI Fund for the Promotion of Joint International Research (International Leading Research) (JP23K20041, to G2P-Japan Consortium and Kei Sato); JSPS KAKENHI Grant-in-Aid for Scientific Research A (JP24H00607, to Kei Sato); Mitsubishi UFJ Financial Group, Inc. Vaccine Development Grant (to Kei Sato); and The Cooperative Research Program (Joint Usage/Research Center program) of Institute for Life and Medical Sciences, Kyoto University (to Kei Sato).

## Declaration of interest

K.S. has consulting fees from Moderna Japan Co., Ltd. and Takeda Pharmaceutical Co. Ltd. and honoraria for lectures from Gilead Sciences, Inc., Moderna Japan Co., Ltd., and Shionogi & Co., Ltd. The other authors declare no competing interests. All authors have submitted the ICMJE Form for Disclosure of Potential Conflicts of Interest. Conflicts that the editors consider relevant to the content of the manuscript have been disclosed.

