## Supplementary Appendix for "Robust antiviral humoral immunity induced by JN.1 monovalent mRNA vaccines against a broad range of SARS-CoV-2 Omicron subvariants including JN.1, KP.3.1.1 and XEC"

**Table of Contents**

**Contents** Page

| | |
|---|---|
| **Materials and Methods** | 2-3 |
| Ethics statement | |
| Human serum collection | |
| Cell culture | |
| Pseudovirus preparation | |
| Neutralization assay | |
| **Table S1.** Human vaccination sera used in this study | 4 |
| **Consortia** | 5-6 |
| **Acknowledgments** | 7 |
| **Supplemental References** | 8 |

### Materials and Methods

#### Ethics statement

All protocols involving specimens from human subjects recruited at Fujiwara Clinic and Namikibashi Clinic were reviewed and approved by the Institutional Review Board of The Institute of Medical Science, The University of Tokyo (approval ID: 2022-29-0915). All human subjects provided written informed consent. All protocols for the use of human specimens were reviewed and approved by the Institutional Review Board of The Institute of Medical Science, The University of Tokyo (approval ID: 2022-29-0915).

#### Human serum collection

JN.1 monovalent mRNA vaccine sera from fully vaccinated individuals who had received Pfizer-BioNTech JN.1 vaccine ("Pfizer-BioNTech" cohort) (two 2-dose vaccinated, two 3-dose vaccinated, five 4-dose vaccinated, three 5-dose vaccinated and three 7-dose vaccinated; fifteen donors, average age: 42.1, range: 27–85, 66.7% male) and those from fully vaccinated individuals who received Daiichi-Sankyo JN.1 vaccine ("Daiichi-Sankyo" cohort) (two 2-dose vaccinated, one 4-dose vaccinated, one 5-dose vaccinated, four 6-dose vaccinated and eleven 7-dose vaccinated; nineteen donors, average age: 57.8, range: 25–84; 26.3% male). For this study, we collected samples before vaccination and three to four weeks (21–29 days) after vaccination. Sera were inactivated at 56°C for 30 minutes and stored at –80°C until use. The details of the convalescent sera are summarized in **Table S1**.

#### Cell culture

The Lenti-X 293T cells (Takara, Cat# 632180) and HOS-ACE2/TMPRSS2 cells (kindly provided by Dr. Kenzo Tokunaga), a derivative of HOS cells (a human osteosarcoma cell line; ATCC CRL-1543) stably expressing human ACE2 and TMPRSS2[1,2] were maintained in Dulbecco's modified Eagle's medium (DMEM) (high glucose) (Wako, Cat# 044- 29765) containing 10% fetal bovine serum (Sigma-Aldrich Cat# 172012-500ML), 100 units penicillin and 100 ug/ml streptomycin (Sigma-Aldrich, Cat# P4333-100ML).

#### Pseudovirus preparation

Plasmids expressing the SARS-CoV-2 spike (S) proteins of B.1.1, BA.5, XBB.1.5, JN.1, KP.2, KP.3, KP.3.1.1 and XEC were prepared in our previous studies.[3-10] Pseudoviruses were prepared as previously described.[6-10] Briefly, lentivirus (HIV-1)-based, luciferase-expressing reporter viruses were pseudotyped with the SARS-CoV-2 S. One prior day of transfection, the LentiX-293T cells were seeded at a density of 2 × $10^6$ cells. The LentiX-293T cells were cotransfected with 1 µg psPAX2-IN/HiBiT (a packaging plasmid encoding the HiBiT-tag-fused integrase[1], 1 µg pWPI-Luc2 (a reporter plasmid encoding a firefly luciferase gene[11] and 500 ng plasmids expressing parental S or its derivatives using TransIT-293 transfection reagent (Mirus, Cat# MIR2704) according to the manufacturer's protocol. Two days post transfection, the culture supernatants were harvested and filtrated. The amount of produced pseudovirus particles was quantified by the HiBiT assay using Nano Glo HiBiT lytic detection system (Promega, Cat# N3040) as previously described.[11] In this system, HiBiT peptide is produced with HIV-1 integrase and forms NanoLuc luciferase with LgBiT, which is supplemented with substrates. At two days postinfection, the infected cells were lysed with a Bright-Glo luciferase assay system (Promega, Cat# E2620), and the luminescent signal produced by firefly luciferase reaction was measured using a GloMax explorer multimode microplate reader 3500 (Promega). The pseudoviruses were harvested and stored at –80°C until use after filtration.

### Neutralization assay

Neutralization assays were performed previously described[6-10] and mainly conducted by a semi-automated high-throughput method using Fluent780 (Tecan).[7-10,12] The SARS-CoV-2 spike pseudoviruses (counting ~100,000 relative light units) and serially diluted (40-fold to 29,160-fold dilution at the final concentration) heat-inactivated sera were manually prepared in a 2-ml 96-well plate (Greiner, Cat# 780271) and in 96-well microplates (ThermoFisher Scientific, Cat# 168136), respectively. The pseudoviruses were dispensed and mixed with the sera in 384-well plates (ThermoFisher Scientific, Cat# 164610) on Fluent780 (Tecan). Pseudoviruses without sera were included as controls. After incubation at 37°C for 1 hour, HOS-ACE2/TMPRSS2 cells (3,000 cells/30 µl) were added to the 20 µl mixture of pseudovirus and serum in the 384-well white plate on the device. Two days post infection, the infected cells were lysed with a Bright-Glo luciferase assay system (Promega, Cat# E2620) on Fluent780 (Tecan), and the luminescent signal was measured and processed using an Infinite200 and a Magellan (Tecan). The assay of each serum sample was performed in quadruplicate, and the 50% neutralization titer ($NT_{50}$) was calculated using Prism 9 (GraphPad Software).

**Table S1. Human vaccination sera used in this study**

| JN.1 vaccine manufacturers | Donor ID | Sex | Age | Date of 1st vaccination (YYYY-MM-DD) | Date of 2nd vaccination (YYYY-MM-DD) | Date of 3rd vaccination (YYYY-MM-DD) | Date of 4th vaccination (YYYY-MM-DD) | Date of 5th vaccination (YYYY-MM-DD) | Date of 6th vaccination (YYYY-MM-DD) | Date of 7th vaccination (YYYY-MM-DD) | Date of sampling (before vaccination) (YYYY-MM-DD) | Date of JN.1 vaccination (YYYY-MM-DD) | Date of sampling (after vaccination) (YYYY-MM-DD) | Time interval between vaccination and the second sampling | Prior infection? (YYYY-MM-DD) |
|---|---|---|---|---|---|---|---|---|---|---|---|---|---|---|---|
| Pfizer/BioNTech | 730011 | Male | 40 | NA | NA | NA | NA | NA | NA | 2023-10-12 | 2024-10-07 | 2024-10-07 | 2024-10-28 | 21 | No |
| Pfizer/BioNTech | 730012 | Female | 85 | NA | NA | NA | NA | NA | NA | 2023-10-05 | 2024-10-07 | 2024-10-07 | 2024-11-01 | 25 | No |
| Pfizer/BioNTech | 730013 | Female | 70 | NA | NA | NA | NA | NA | NA | 2023-11-16 | 2024-10-07 | 2024-10-07 | 2024-11-05 | 29 | No |
| Pfizer/BioNTech | UT0001 | Male | 35 | 2021-07-19M | 2021-08-20M | 2022-05-13M | 2022-11-22MBA.4/5 | 2023-10-01MXBB | - | - | 2024-10-02 | 2024-10-03 | 2024-10-29 | 26 | Yes (2023-07-03) |
| Pfizer/BioNTech | UT0002 | Female | 55 | 2021-06-26P | 2021-07-17P | 2022-02-05P | 2022-11-16PBA.4/5 | - | - | - | 2024-09-27 | 2024-10-04 | 2024-10-29 | 25 | No |
| Pfizer/BioNTech | UT0003 | Male | 29 | NA | 2023-2-15MBA.4/5 | - | - | - | - | - | 2024-10-02 | 2024-10-03 | 2024-10-29 | 26 | No |
| Pfizer/BioNTech | UT0004 | Male | 27 | 2021-07-27M | NA | 2022-07-10P | 2023-01-18PBA.4/5 | - | - | - | 2024-10-02 | 2024-10-03 | 2024-10-29 | 26 | No |
| Pfizer/BioNTech | UT0005 | Male | 42 | 2021-06-17P | 2021-07-07P | 2022-03-28M | 2022-10-27MBA.4/5 | 2023-09-20PXBB | - | - | 2024-10-02 | 2024-10-03 | 2024-10-25 | 22 | Yes (2023-06-29) |
| Pfizer/BioNTech | UT0006 | Female | 33 | 2021-03-03P | 2021-8-5P | 2022-2P | 2022-11P | 2023-11P | - | - | 2024-09-27 | 2024-10-04 | 2024-10-29 | 25 | Yes (2021-06) |
| Pfizer/BioNTech | UT0007 | Male | 31 | 2021-07-28M | 2021-08-25M | 2022-03-10M | - | - | - | - | 2024-10-02 | 2024-10-03 | 2024-10-29 | 26 | Yes (2022-08-07) |
| Pfizer/BioNTech | UT0008 | Male | 27 | 2021-08-06M | 2021-09-03M | - | - | - | - | - | 2024-10-02 | 2024-10-03 | 2024-10-29 | 26 | Yes (2022-11-05) |
| Pfizer/BioNTech | UT0009 | Female | 54 | 2021-08-18P | 2021-09-08P | 2022-04-13P | 2022-10-21P | - | - | - | 2024-10-02 | 2024-10-04 | 2024-10-29 | 25 | Yes (2023-07-24) |
| Pfizer/BioNTech | UT0010 | Male | 40 | 2021-06-28 | 2021-07-19 | 2022-05-17 | 2022-09-03 | - | - | - | 2024-10-02 | 2024-10-03 | 2024-10-29 | 26 | No |
| Pfizer/BioNTech | UT0011 | Male | 29 | 2021-05-28S | 2021-06-30S | 2021-12-21J | - | - | - | - | 2024-10-02 | 2024-10-03 | 2024-10-29 | 26 | Yes (2023-05) |
| Pfizer/BioNTech | UT0012 | Male | 35 | 2021-08-27M | 2021-09-24M | 2022-04-16M | 2023-09-29PXBB | - | - | - | 2024-10-02 | 2024-10-03 | 2024-10-29 | 26 | Yes (2023-08-03) |
| Daiichi-Sankyo | F000100 | Male | 45 | NA | NA | NA | NA | NA | 2023-10-27 | - | 2024-10-01 | 2024-10-01 | 2024-10-29 | 28 | No |
| Daiichi-Sankyo | F079661 | Male | 62 | NA | NA | NA | 2022-10 | - | - | - | 2024-10-01 | 2024-10-01 | 2024-10-25 | 24 | No |
| Daiichi-Sankyo | F037838 | Female | 69 | NA | NA | NA | NA | NA | NA | 2023-11-14 | 2024-10-01 | 2024-10-01 | 2024-10-22 | 21 | No |
| Daiichi-Sankyo | F075080 | Female | 74 | NA | NA | NA | NA | NA | NA | 2023-09-29 | 2024-10-01 | 2024-10-01 | 2024-10-22 | 21 | No |
| Daiichi-Sankyo | F072023 | Female | 57 | NA | NA | NA | NA | NA | NA | 2023-11-10 | 2024-10-01 | 2024-10-01 | 2024-10-22 | 21 | No |
| Daiichi-Sankyo | F055336 | Female | 75 | NA | NA | NA | NA | NA | NA | 2023-10-06 | 2024-10-02 | 2024-10-02 | 2024-10-25 | 23 | No |
| Daiichi-Sankyo | F018678 | Female | 63 | NA | NA | NA | NA | NA | NA | 2023-09-29 | 2024-10-02 | 2024-10-02 | 2024-10-23 | 21 | Yes (2023-03, 2024-08) |
| Daiichi-Sankyo | F072088 | Female | 71 | NA | NA | NA | NA | NA | NA | 2023-09-26 | 2024-10-03 | 2024-10-03 | 2024-10-30 | 27 | No |
| Daiichi-Sankyo | F035951 | Female | 74 | NA | NA | NA | NA | NA | 2023-06-13 | - | 2024-10-04 | 2024/10/04 | 2024-10-25 | 21 | No |
| Daiichi-Sankyo | F035846 | Male | 82 | NA | NA | NA | NA | NA | NA | 2023-09-26 | 2024-10-04 | 2024-10-04 | 2024-10-25 | 21 | No |
| Daiichi-Sankyo | F050351 | Female | 79 | NA | NA | NA | NA | NA | NA | 2023-09-26 | 2024-10-04 | 2024-10-04 | 2024-10-29 | 25 | No |
| Daiichi-Sankyo | F000300 | Male | 62 | NA | NA | NA | NA | 2023-11-21 | - | - | 2024-10-04 | 2024-10-04 | 2024-10-29 | 25 | Yes (2023-07) |
| Daiichi-Sankyo | F079996 | Male | 84 | NA | NA | NA | NA | NA | 2023-10-24 | - | 2024-10-04 | 2024-10-04 | 2024-10-25 | 21 | Yes (2022-10) |
| Daiichi-Sankyo | F000200 | Female | 45 | NA | NA | NA | NA | NA | 2023-10-28 | - | 2024-10-03 | 2024-10-03 | 2024-10-24 | 21 | No |
| Daiichi-Sankyo | 730016 | Female | 36 | NA | 2021 | - | - | - | - | - | 2024-10-10 | 2024-10-10 | 2024-10-31 | 21 | Yes (2022-12) |
| Daiichi-Sankyo | 730017 | Female | 28 | NA | NA | NA | NA | NA | NA | 2023-10 | 2024-10-10 | 2024-10-10 | 2024-10-31 | 21 | Yes (2022-09) |
| Daiichi-Sankyo | 730018 | Female | 40 | NA | 2021 | - | - | - | - | - | 2024-10-10 | 2024-10-10 | 2024-10-31 | 21 | Yes (2021-08) |
| Daiichi-Sankyo | 730020 | Female | 25 | NA | NA | NA | NA | NA | NA | 2023-08 | 2024-10-10 | 2024-10-10 | 2024-10-31 | 21 | Yes (2023-08) |
| Daiichi-Sankyo | 730019 | Female | 28 | NA | NA | NA | NA | NA | NA | 2023-10 | 2024-10-10 | 2024-10-10 | 2024-10-31 | 21 | No |

NA, not applicable.

A, Astrazeneca; P, Pfizer/BioNTech; M, Moderna; S, Sinovac; J, Jenner

BA1/2, BA.1/2 bivalent vaccine; BA4/5, BA.4/5 bivalent vaccine; XBB, XBB.1.5 monovalent vaccine

**Consortia**

**The Genotype to Phenotype Japan (G2P-Japan) Consortium**

**The Institute of Medical Science, The University of Tokyo, Japan**
Jumpei Ito, Naoko Misawa, Arnon Plianchaisuk, Ziyi Guo, Alfredo Hinay Jr., Kaoru Usui, Wilaiporn Saikruang, Spyros Lytras, Shusuke Kawakubo, Luca Nishimura, Yusuke Kosugi, Shigeru Fujita, Luo Chen, Jarel Elgin M. Tolentino, Lin Pan, Wenye Li, Maximilian Stanley Yo, Daniel Arnold, Yukun Zhu, Mai Suganami, Mika Chiba, Keiko Iida, Naomi Ohsumi, Shiho Tanaka, Eiko Ogawa, Kyoko Yasuda, Kaho Okumura, Tsuki Fukuda, Tamaki Yoshihara, Keiko Koizumi, Hiroaki Unno

**Hokkaido University, Japan**
Takasuke Fukuhara, Tomokazu Tamura, Rigel Suzuki, Saori Suzuki, Shuhei Tsujino, Hayato Ito, Hirofumi Sawa, Naganori Nao, Keita Matsuno, Keita Mizuma, Jingshu Li, Izumi Kida, Yume Mimura, Yuma Ohari, Shinya Tanaka, Masumi Tsuda, Lei Wang, Yoshikata Oda, Zannatul Ferdous, Kenji Shishido, Hiromi Mohri, Miki Iida

**Tokyo Metropolitan Institute of Public Health**
Kenji Sadamasu, Kazuhisa Yoshimura, Hiroyuki Asakura, Isao Yoshida, Mami Nagashima

**Tokai University, Japan**
So Nakagawa

**Kyoto University, Japan**
Kotaro Shirakawa, Akifumi Takaori-Kondo, Kazuo Takayama, Rina Hashimoto, Sayaka Deguchi, Yukio Watanabe, Yoshitaka Nakata, Hiroki Futatsusako, Ayaka Sakamoto, Naoko Yasuhara, Takao Hashiguchi, Tateki Suzuki, Kanako Kimura, Jiei Sasaki, Yukari Nakajima, Hisano Yajima

**Hiroshima University, Japan**
Takashi Irie, Ryoko Kawabata

**Kyushu University, Japan**
Kaori Tabata

**Kumamoto University, Japan**
Terumasa Ikeda, Hesham Nasser, Ryo Shimizu, MST Monira Begum, Michael Jonathan, Yuka Mugita, Sharee Leong, Otowa Takahashi, Takamasa Ueno, Chihiro Motozono, Mako Toyoda

**University of Miyazaki, Japan**
Akatsuki Saito, Anon Kosaka, Miki Kawano, Natsumi Matsubara, Tomoko Nishiuchi

**Charles University, Czechia**
Jiri Zahradnik, Prokopios, Andrikopoulos, Miguel Padilla-Blanco, Aditi Konar, Ruojin Tuan

### Acknowledgments

We would like to thank all members of The Genotype to Phenotype Japan (G2P-Japan) Consortium. We thank Kenzo Tokunaga (National Institute of Infectious Diseases, Japan) for sharing materials and Mika Chiba, Kyoko Yasuda, Keiko Iida, Tsuki Fukuda, Tamaki Yoshihara and Keiko Koizumi (Division of Systems Virology, University of Tokyo, Japan) for supporting experimental assays.
